# A Dynamics-Informed Machine Learning Framework for Inhibitor Optimization Against Quickly Evolving Targets to Avoid Resistance

**DOI:** 10.64898/2026.09.25.754496

**Authors:** Lauren E. Intravaia, Jagroop Kaur, Ala M. Shaqra, Mina Henes, Klajdi Kosovrasti, Viviana Schroeder, Ellen A. Nalivaika, Akbar Ali, Nese Kurt Yilmaz, Celia A. Schiffer

## Abstract

Drug resistance often emerges through mutations that reshape protein conformational ensembles, allowing inhibitor potency to be altered without directly disrupting protein-ligand contacts. Consequently, the molecular mechanisms underlying these effects remain difficult to identify, hindering the design of inhibitors with durable potency against rapidly evolving drug targets. To address this challenge, we developed ROBUST, a physics-informed machine learning framework that integrates experimental measurements, molecular dynamics simulations, and interpretable statistical modeling to identify energetic determinants of inhibitor potency. Using HIV-1 protease as a stringent model system for drug resistance, we determined 157 new inhibition constants and integrated them with prior measurements to establish a comprehensive experimental dataset of 364 protease-inhibitor pairs. Analysis of the corresponding molecular dynamics simulations revealed a compact network of energetic interactions that quantitatively explains how resistance mutations and inhibitor modifications jointly reshape molecular recognition and inhibitor potency across the multidimensional resistance landscape. Models constructed from these energetic signatures predicted binding free energies with high accuracy (RMSE < 1 kcal mol^−1^) and successfully generalized to unseen darunavir analogs. Importantly, ROBUST accurately prioritized unseen analogs based on their potency across the variant set, demonstrating its utility for identifying compounds with improved resistance profiles. Together, these results uncover the energetic principles through which resistance mutations and inhibitor modifications alter potency and establish ROBUST as a mechanistically interpretable framework for translating these insights into resistance-aware inhibitor optimization against rapidly evolving drug targets.

## INTRODUCTION

Drug resistance remains a major barrier in developing long-lasting therapeutics. Despite the rapid design, development, and deployment of new drugs for infectious diseases and cancers, resistance can emerge and ultimately render effective treatments obsolete. This imposes substantial economic and clinical burdens which are projected to worsen in the coming decades.^1, 2^ Resistance has been documented across diverse drug targets, including viral proteases^3–7^, reverse transcriptases^8, 9^, polymerases^10–13^, and integrases^14, 15^; bacterial targets such as dihydrofolate reductases^16–19^; and cancer targets including multiple kinases^20–26^. Consequently, strategies that anticipate and mitigate target-based resistance early in drug development are urgently needed. Here, we use HIV-1 protease as a stringent model of resistance evolution to identify fundamental principles for optimizing inhibitors against rapidly evolving targets.

Lead optimization is a critical stage of drug development in which small molecules are systematically modified to improve drug-like properties, including potency^27^. When a three-dimensional molecular structure of the target-ligand complex is available, potency can be enhanced through chemical modifications predicted to improve binding interactions within the complex^28, 29^. To explore the effect of broader chemical modifications, many structure-based lead optimization campaigns rely on computational methods to estimate inhibition potency and prioritize analogs for synthesis and testing^30–32^. Among physics-based approaches, alchemical free energy perturbation (FEP) calculations remain the most established method for predicting potency. However, their application becomes more challenging when mutations induce substantial conformational reorganization that alters the ensembles governing inhibitor binding^33^. Moreover, while FEP is broadly transferable across targets, it is not intended to identify the energetic interaction networks that mechanistically govern inhibition and resistance.

Machine learning (ML) has therefore emerged as a promising approach for potency prediction and resistance modeling. Broadly transferable foundation models, such as Boltz-2, have advanced binding affinity estimation, yet their application to resistant targets remains limited by incomplete representation of the conformational and energetic features that govern inhibitor resistance^34^. At the same time, target-specific ML methods for lead optimization and resistance prediction have shown considerable promise^35–55^, but many rely on geometric descriptors, nonlinear ML algorithms, or classification schemes that do not directly reveal the physical interactions governing inhibition. Consequently, a central unmet need is a physically interpretable framework that not only predicts potency but also identifies the energetic interaction networks that maintain potency in the face of evolutionary pressure, providing a mechanistic basis for prioritizing chemical modifications during lead optimization.

In prior work, we demonstrated that distal resistance mutations perturb the conformational ensemble of HIV-1 protease and modulate inhibitor potency through changes in protein dynamics rather than direct disruption of binding contacts^56–59^. We subsequently showed that molecular dynamics (MD)-derived energetic descriptors can be incorporated into interpretable ML models that predict inhibitor resistance and identify mechanistic determinants of potency loss^60, 61^. These studies established that resistance-associated changes in protein dynamics can be captured through a small set of recurring energetic interactions. However, it remains unclear whether such energetic signatures are sufficiently robust to generalize across extensive chemical variation and guide resistance-aware lead optimization. We hypothesized that inhibitor potency is governed by conserved dynamic features of the conformational ensemble that are preserved across mutational and chemical variation. To test this hypothesis, we developed ROBUST, a parallel MD-coupled ML framework that extracts recurring energetic descriptors from protein-ligand ensembles and evaluates their ability to predict experimentally measured potencies. Using HIV-1 protease as a stringent model of resistance evolution, we show that a small number of dynamics-derived energetic interactions quantitatively capture potency across resistant variants and chemically distinct inhibitor analogs. More broadly, these studies establish a framework for identifying resistance-aware molecular design rules that connect conformational dynamics, inhibitor chemistry, and potency.

## RESULTS

### A Multidimensional Landscape of HIV-1 Protease Resistance

To map the HIV-1 protease resistance landscape, we determined 157 new inhibition constants spanning diverse protease variants and structurally distinct darunavir analogs. These measurements were combined with 207 previously reported values^62, 63^ to establish a dataset comprising 364 protease variant-inhibitor pairs spanning extensive mutational and chemical diversity (Figure 1; Table S1).

**Figure 1.**
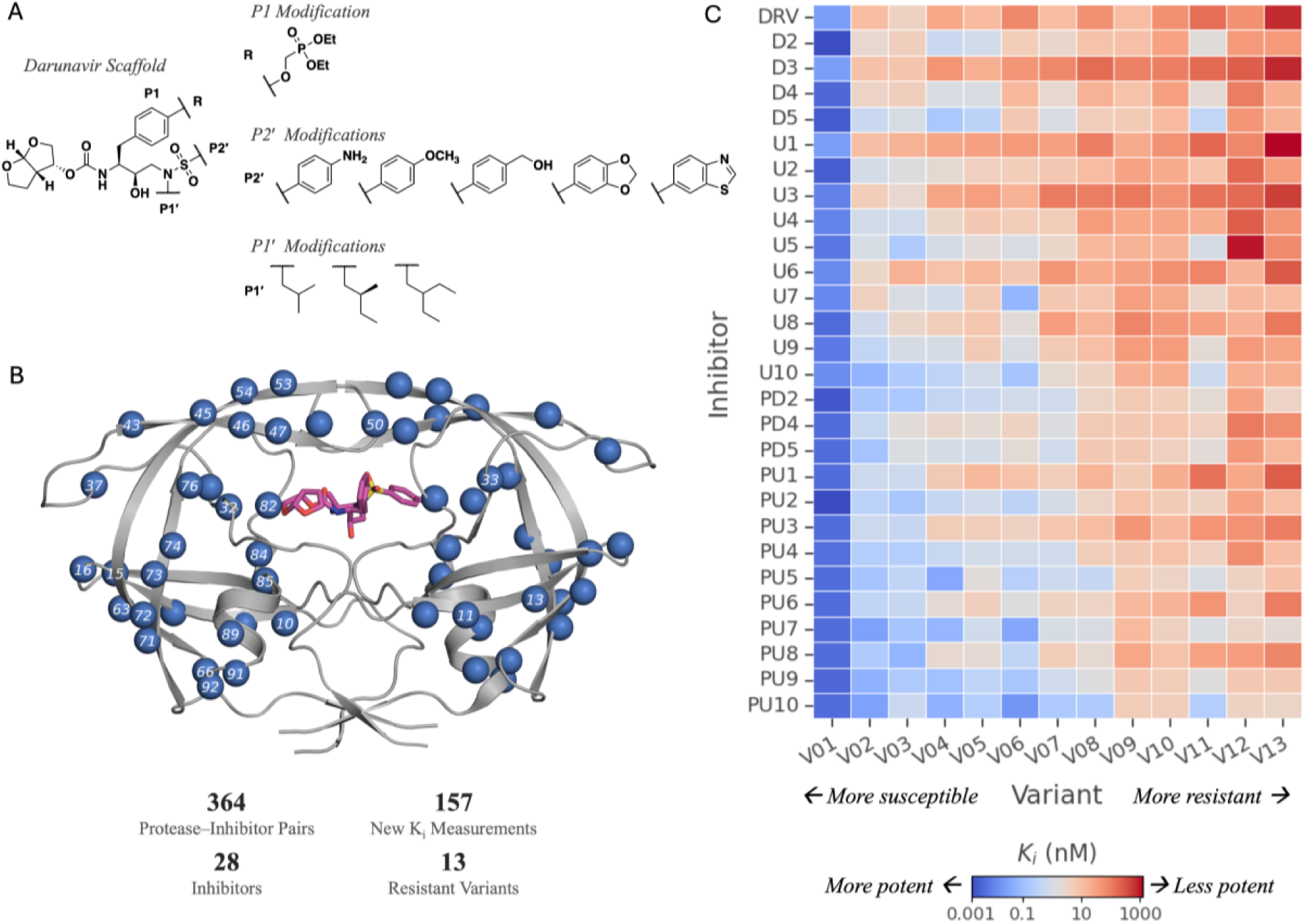
Experimental landscape of mutational and chemical diversity in HIV-1 protease inhibition. (A) Chemical structure of darunavir scaffold and the modifications in the 27 inhibitor analogs. (B) Crystal structure of HIV-1 protease bound to darunavir (magenta sticks) in cartoon representation (PDB ID: 6DGX) where amino acid residues with mutations in the dataset are indicated by blue spheres and residue numbers. (C) Experimental inhibition constants for all 364 protease variant-inhibitor pairs colored blue to red for increasing resistance.

The inhibitor set includes darunavir and 27 analogs that contain modifications at the P1, P1′, and P2′ moieties (binding to the corresponding S1, S1′ and S2′ subsites of the active site) (Figure 1A; for the complete inhibitor chemical structures see Figure S1). The panel comprises 13 HIV-1 protease variants with substitutions at 28 of the 99 amino acid residues, including mutations both within and distal to the inhibitor binding site (Figure 1B; the corresponding mutation profiles are summarized in Table S2). The individual HIV-1 protease variants have between one and fourteen substitutions relative to a wild-type sequence (the wild-type variant was not included as the sub-picomolar affinity could not be accurately measured). Collectively, the experimental *K_i_* values span picomolar- to micromolar-scale potency (Figure 1C) and reveal distinct resistance profiles across both protease variants and darunavir analogs (Figure S2 and S3). By systematically sampling both protease sequence and inhibitor chemistry, this large-scale experimental dataset provides a comprehensive map of the multidimensional HIV-1 protease resistance landscape and enables direct interrogation of the molecular determinants of inhibitor potency and resistance.

### Resistance Mutations and Inhibitor Modifications Reshape a Shared Energetic Networ

We sought to determine whether the complex resistance landscape of HIV-1 protease could be reduced to a compact set of physically meaningful energetic interactions. Toward this goal, pairwise residue-residue and residue-inhibitor interaction energies were extracted from equilibrium MD trajectories and used to construct a comprehensive physics-based descriptor space comprising 1,503 van der Waals (vdW) interactions and 985 electrostatics (ES) interactions for each complex. Repeated feature-selection analyses were then performed on the full dataset to identify energetic interactions that consistently linked conformational dynamics to experimental potency across both variant sequence and inhibitor chemistry.

Despite the complexity of the energetic descriptor space, inhibitor potency was associated with a remarkably small set of recurring interactions. Across the complete mutational and chemical landscape represented in the dataset, three descriptors occurred in the majority of Akaike Information Criterion (AIC)-supported models (Figure 2A; see also Methods). Mapping the recurring energetic descriptors onto the protease structure revealed that the dominant interactions cluster within a flap-centered communication network linking inhibitor recognition and flap dynamics (Figure 2B): The vdW interaction 10B–82B reports on active-site packing near the inhibitor-binding pocket whereas vdW 47B–LIG captures interactions between the flexible flap region and the inhibitor P2′ moiety. The ES interaction 46B–54B reflects stabilization of the flap β-sheet architecture that regulates conformational accessibility of the active site. Importantly, the three dominant descriptors individually correlated with experimental potency (Figure 2C), indicating that these energetic interactions report on distinct yet interconnected aspects of the resistance-modulated binding landscape. Taken together, these observations suggest that a limited number of energetically coupled descriptors capture the dominant determinants of inhibitor potency, providing a compact representation of the resistance landscape suitable for quantitative modeling.

**Figure 2.**
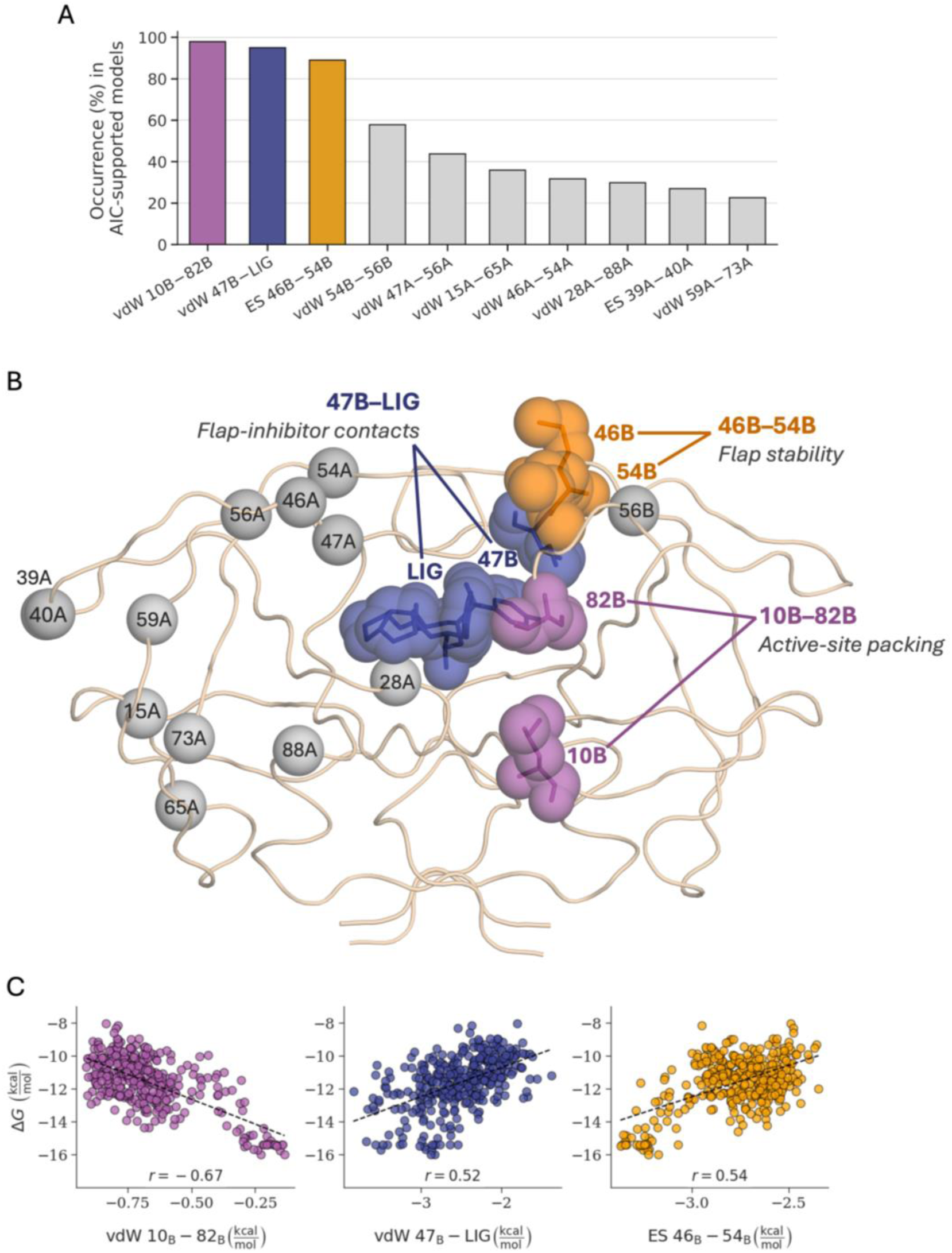
A compact energetic interaction network governing inhibitor potency and resistance. (A) Frequency of occurrence of recurring features among AIC-supported models generated from 100 independent train/test partitions, and (B) mapping of these onto the HIV-1 protease structure. The three dominant interactions are highlighted in color. (C) Correlation of the three dominant descriptors with experimental binding free energy.

### Energetic Signatures Quantitatively Explain Inhibitor Potency and Resistance

The recurring descriptors did not behave independently but instead formed an interconnected energetic network, suggesting that resistance emerges from coordinated redistribution of interactions rather than disruption of a single contact (Figure 3A). To determine whether this dynamics-derived energetic signature could quantitatively reproduce experimental potency, we evaluated predictive models constructed from the most frequently selected descriptors. Increasing the number of descriptors improved predictive performance, with model accuracy plateauing at approximately five to six features (Figure 3B), indicating that a compact feature set captures most of the predictive information contained within the broader energetic network. Consistent with this observation, a model incorporating vdW interactions 10B–82B, 47B–LIG, 54B–56B, and 47A– 56A together with the ES interaction 46B–54B accurately reproduced the experimental potency landscape, achieving strong cross-validated performance (RMSE = 0.88 kcal/mol; Pearson *r* = 0.85) and comparable test-set performance (RMSE = 0.81 kcal/mol; Pearson *r* = 0.90; Figure 3C). Across the 100 random train/test splits, the resulting models consistently achieved high Pearson correlations, low prediction errors, and stable coefficients of determination (Figure 3D), demonstrating the robustness of the identified energetic signature. Together, these results indicate that a small set of energetically coupled interactions is sufficient to describe potency across resistant HIV-1 protease variants and chemically diverse darunavir analogs. Importantly, although three interactions emerged as the dominant mechanistic descriptors, the inclusion of additional correlated features improved quantitative prediction, consistent with a compact but partially redundant energetic network governing inhibitor potency.

**Figure 3.**
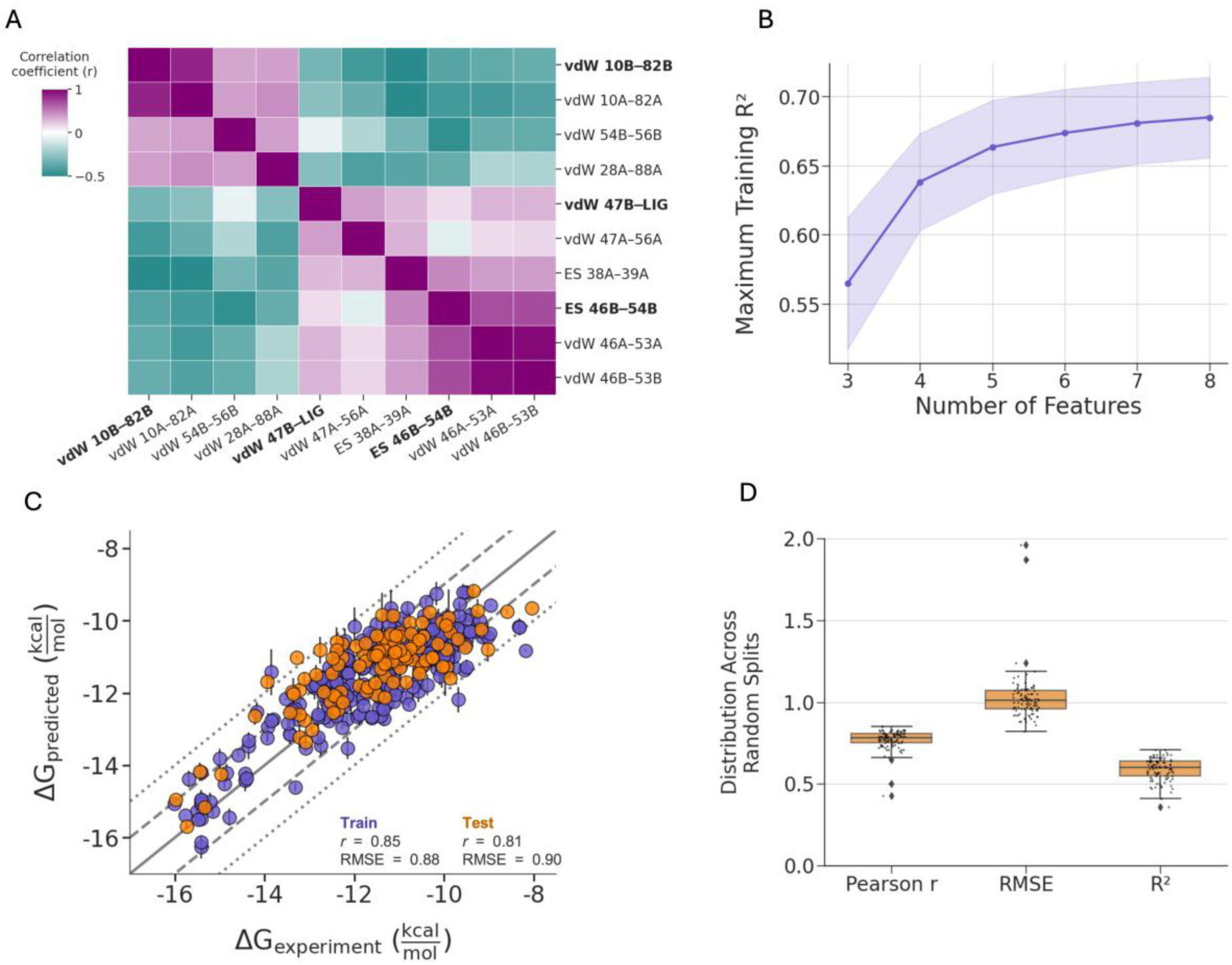
Models constructed from the energetic signatures reproduce inhibitor potency across the resistance landscape. (A) Intercorrelation matrix of the ten most recurrent energetic descriptors identified across AIC-supported models. The three most frequent features, vdW 10B–82B, vdW 47B–LIG, and ES 46B–54B, are indicated in bold. Positive and negative correlations are shown in magenta and teal, respectively. (B) Maximum training set R^2^ achieved by AIC-supported models as a function of feature number across 100 random train/test splits. The shaded region indicates the standard deviation across splits. (C) Experimental versus predicted binding affinity for a representative ordinary-least squares regression model constructed from the recurring energetic descriptors. Training and test data are shown in purple and orange, respectively. Solid, dashed, and dotted lines indicate the identity line and deviations of ±1 and ±2 kcal/mol. (D) Distribution of test-set performance metrics across 100 random train/test splits, including Pearson correlation coefficients, root-mean-square errors (RMSE), and coefficients of determination (R^2^).

### Molecular Determinants of Potency Generalize to Unseen Inhibitor Modifications

The critical test of any lead-optimization framework is whether it can successfully predict compounds that lie outside the chemistry used for model development. To evaluate whether the energetic signatures identified by ROUBST generalize beyond the training chemical space, we generated three independent train/test partitions in which inhibitors containing specific P1, P1′, or P2′ substituents were excluded from model training and reserved for external validation (Figure 4A–C). Despite these stringent holdout conditions, compact energetic signatures retained strong predictive power across all three validation schemes (Figure S5A–F). For the P1 holdout, a linear regression model constructed from a subset of energetic descriptors (Figure S5A,B) accurately predicted the potency of inhibitors containing the phosphonate substitution excluded from model training (RMSE = 1.08 kcal/mol; Pearson *r* = 0.79; Figure 4D). Similarly, the P1′ holdout model successfully predicted inhibitors containing the bulkier tert-butyl substituent (RMSE = 0.89 kcal/mol, Pearson r = 0.84; Figure 4E), whereas the P2′ holdout model generalized to inhibitors containing cyclic thiazole or dioxolane substituents (RMSE = 0.98 kcal/mol, Pearson’s r = 0.80; Figure 4F). Across all three validation schemes, 64-72% of test complexes were predicted within 1 kcal/mol of their experimental binding free energy (Figure S6B-D). Notably, the predictive performance for unseen chemotypes was comparable to that obtained for randomly partitioned datasets, indicating that the identified energetic descriptors capture transferable structure-dynamics-potency relationships rather than features specific to individual inhibitor modifications. Thus, the mutational and chemical diversity converged on a common energetic interaction network that remains predictive beyond the chemical space used for model development.

**Figure 4.**
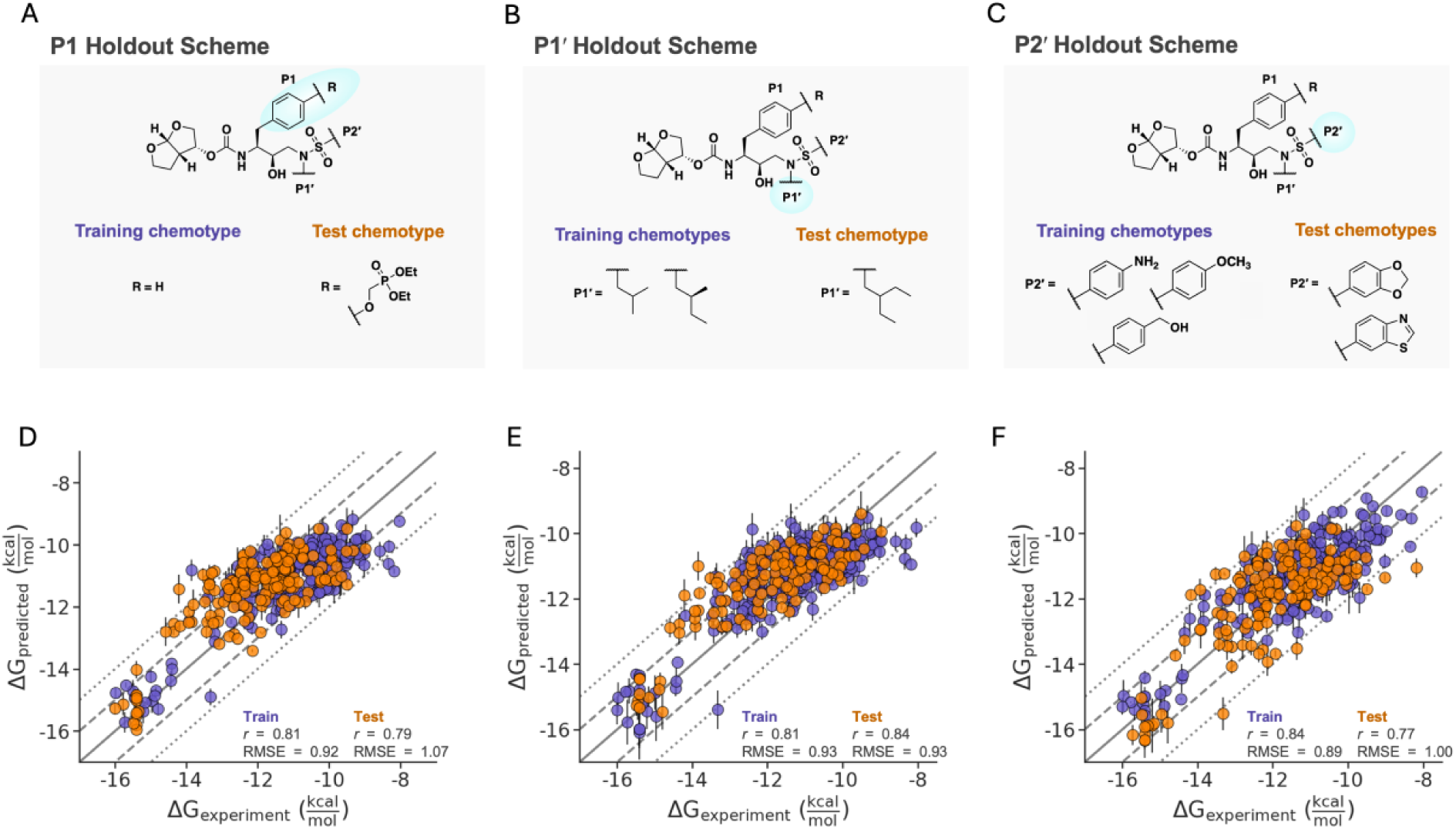
Dynamics-derived energetic signatures generalize beyond the chemical training space. Training and test sets were constructed by holding out inhibitors containing specific (A) P1, (B) P1′ or (C) P2′ substituents from model training. Training and held-out chemotypes are shown in orange and purple, respectively. Experimental versus predicted binding free energies for the corresponding holdout models are shown in panels (D), (E), and (F). Training and held-out test compounds are shown in purple and orange, respectively. Solid, dashed, and dotted lines indicate the identity line and deviations of ±1 and ±2 kcal/mol. Performance statistics for training and test sets are reported within each panel.

### ROBUST Prioritizes Inhibitors with Improved Resistance Profiles

Predicting potency is valuable, but the ultimate objective of lead optimization during drug discovery is to identify which analogs should be prioritized for synthesis and experimental evaluation. To assess the utility of ROBUST as a resistance-aware prioritization tool, we averaged the predicted potencies of each inhibitor against the panel of 13 variants for the P1, P1′, and P2′ holdout datasets. In each case, the models correctly identified inhibitor modifications with improved average potency relative to darunavir, with only a small number of exceptions (Figure 5A-C). Importantly, the held-out inhibitors were not included during model training, providing a stringent test of prospective ranking performance.

**Figure 5.**
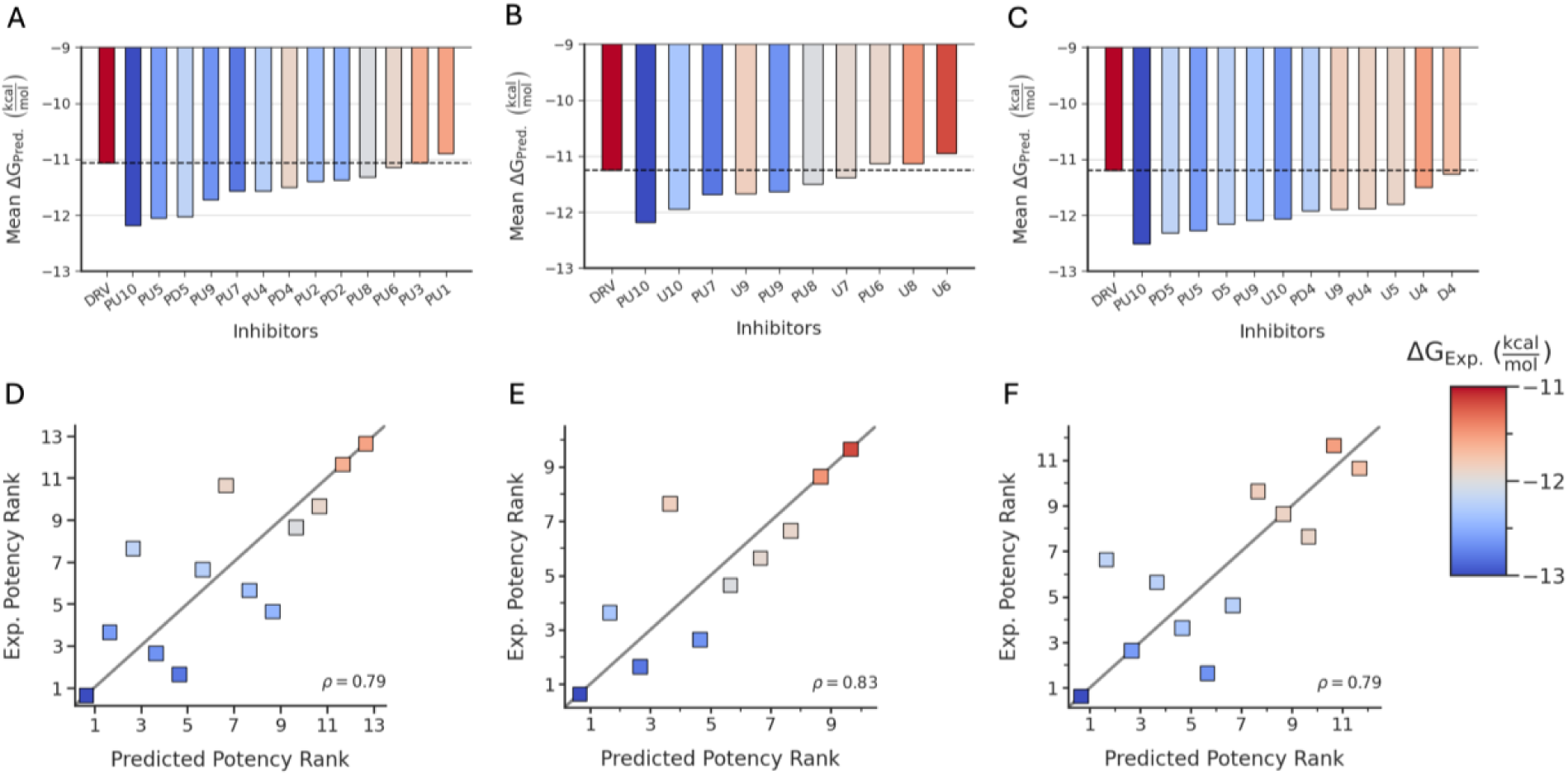
A compact energetic interaction network enables prioritization of unseen analogs across resistant variants. For the (A, D) P1, (B, E) P1′ and (C, F) P2′ holdout datasets, inhibitors were ranked according to predicted mean binding free energies against the panel of resistant HIV-1 protease variants. Panels A-C show the predicted mean potencies of the held-out inhibitor relative to DRV (dashed line), with bars colored according to experimentally determined mean binding free energies. Panels D-F compare predicted and experimental potency rankings for the corresponding compounds, with points colored by the same experimental mean binding free energies.

To evaluate prioritization accuracy, inhibitors were ranked according to their predicted average potency across the variant panel and compared with the corresponding experimental rankings. The predicted rankings closely reproduced the experimental ordering of the unseen analogs for all three holdout schemes (P1: ρ = 0.79; P1′: ρ = 0.83; P2′: ρ = 0.79; Figure 5D-F). These results demonstrate that dynamics-derived energetic signatures identified by ROBUST retain predictive value beyond individual potency estimates and can successfully prioritize compounds according to their breadth of activity against resistant variants. This result suggests that the ROBUST approach can be used not only to explain potency trends and predict unseen chemotypes but also to guide resistance-aware lead optimization by identifying analogs with the greatest potential for broad inhibition of resistant HIV-1 protease variants.

## DISCUSSION

Despite decades of investigation into HIV-1 protease drug resistance, it has remained difficult to determine how resistance mutations and inhibitor modifications jointly shape inhibitor potency. Remarkably, despite extensive sequence and chemical diversity, inhibitor potency was governed by a compact network of recurring energetic interactions that consistently mapped to residues involved in regulating HIV-1 protease flap architecture and active-site accessibility. Although three energetic descriptors emerged most frequently during feature selection, predictive performance was enhanced by incorporating several additional correlated interactions. Consistent with the ensemble nature of protein dynamics, multiple interactions appear to report on related underlying conformational processes, allowing distinct descriptor subsets to capture similar information about the resistance landscape. These findings support a view of resistance as a network phenomenon, in which potency is modulated through redistribution of energetic communication pathways rather than disruption of individual protein-inhibitor contacts. The ROBUST framework explains how distal mutations and inhibitor modifications can substantially alter potency despite producing only modest changes in static protein-inhibitor structures. Consequently, resistance mutations and inhibitor modifications appear to act through a common dynamic control mechanism, with potency determined by the extent to which favorable interactions within this network are preserved.

The breadth of the resistance landscape examined here, encompassing 364 protease-inhibitor pairs across extensive sequence and chemical diversity, enabled identification of energetic signatures that remained robust across multiple mutational and chemical contexts. Importantly, this network-level representation generalized successfully to unseen darunavir modifications. When these chemotypes were excluded from model construction, the resulting models retained predictive power and accurately prioritized analogs according to their potency across the variant panel. Successful transfer across multiple classes of chemical modification demonstrates that ROBUST identifies energetic features that remain predictive beyond the chemical space used for model development. The practical implication is that inhibitor optimization may be guided by preservation of these energetic features rather than maximizing potency against a single protease variant. Within this framework, the most promising analogs are those that maintain favorable interactions across many resistant variants, thereby preserving potency in the face of evolution. ROBUST therefore provides a physics-based strategy for prioritizing inhibitor modifications with improved resistance profiles.

ROBUST is intended to complement rather than replace existing computational approaches for lead optimization. Physics-based free-energy calculations remain among the most rigorous methods for estimating binding affinities, while modern ML approaches can achieve strong predictive performance across diverse systems. However, these methods are not explicitly designed to identify the molecular mechanisms that govern durable inhibition against rapidly evolving drug targets. In contrast, ROBUST explicitly links potency predictions to physically interpretable interaction energies derived from protein dynamics. Notably, ROBUST achieved sub-kcal mol^−1^ predictive accuracy despite encompassing substantial mutational and chemical diversity, suggesting that a small set of dynamics-derived energetic descriptors captures fundamental features of the resistance landscape.

More broadly, these findings illustrate how combining experimental measurements, MD simulations, and interpretable ML can transform complex biological datasets into mechanistic models of potency and resistance. While developed using HIV-1 protease, ROBUST suggests a general design principle for rapidly evolving therapeutic targets: durable potency may depend on preserving favorable, dynamics-coupled energetic interaction networks rather than optimizing individual protein-ligand contacts. Similar resistance-associated conformational redistribution has been observed in diverse viral enzymes, kinases, and other rapidly evolving proteins, suggesting that compact energetic interaction networks may represent a broadly useful principle for understanding molecular recognition in evolving systems. In this sense, the recurring signatures identified here constitute a dynamics-derived energetic pharmacophore, capturing conserved energetic relationships across both sequence and chemical diversity. Notably, the specific descriptors selected varied somewhat across training subsets while repeatedly mapping to the same underlying energetic network, suggesting that potency is governed by network-level mechanisms rather than individual contacts. Together, these results uncover energetic principles that link target evolution and inhibitor chemistry to potency and establish ROBUST as a mechanistically interpretable framework for resistance-aware inhibitor design. More broadly, our findings suggest that dynamics-derived energetic interaction networks may represent a generalizable basis for understanding and overcoming resistance in rapidly evolving therapeutic targets.

## EXPERIMENTAL METHODS

### Determination of Experimental Potency (*K_i_* values)

Enzyme inhibition constants (*K_i_* values) were determined using a fluorescence-based HIV-1 protease inhibition assay as previously described^64, 65^. In the present study, 157 previously unreported *K_i_* values were determined for a panel of HIV-1 protease variant-inhibitor pairs. Briefly, inhibitors were serially diluted in 96-well plates, including a no-inhibitor control, and incubated with protease (0.75-5.0 nM) for 1 h. A 10-amino acid substrate containing an optimized protease cleavage site^64^ and an EDANS/DABCYL FRET pair was dissolved in 4% DMSO and added to the reaction mixture. Fluorescence was monitored using a PerkinElmer EnVision plate reader (λ_ex = 340 nm, λ_em = 492 nm) for approximately 60 min. Data was analyzed with GraphPad Prism (version 10), and darunavir was included as a control in all assays.

### Construction of the Experimental Resistance Dataset

The HIV-1 protease variants and darunavir analogs analyzed in this study have been described previously^62, 63, 66, 67^. The experimental resistance dataset was constructed by combining 157 newly determined *K_i_* values with 207 previously reported measurements^62, 63^ yielding a total of 364 protease-inhibitor pairs. The dataset comprises 13 HIV-1 protease variants and 28 inhibitors, including darunavir and 27 analogs, spanning extensive sequence and chemical diversity. Collectively, the measurements encompass inhibition constants ranging from the picomolar to micromolar regime and provide a multidimensional experimental map of the HIV-1 protease resistance used for all subsequent modeling and analysis.

## COMPUTATIONAL METHODS

### Overview of the ROBUST Workflow

The ROBUST workflow integrates experimental potency measurements, MD simulations, energetic feature extraction, and machine learning to identify compact interaction signatures associated with inhibitor potency^60, 61, 68^. Briefly, experimentally characterized protease-inhibitor complexes were simulated by MD. Pairwise residue-residue and residue-inhibitor interaction energies were subsequently extracted from equilibrium trajectories and used as inputs for feature-selection and regression analyses. Recurring descriptors were identified by elastic net regression and incorporated into ordinary least-squares (OLS) models to generate physically interpretable predictors of inhibitor potency. The resulting framework was used to identify energetic signatures of potency, evaluate predictive performance, assess generalization to unseen inhibitor chemotypes, and prioritize compounds for resistance-aware lead optimization.

### Preparation of HIV-1 Protease-Inhibitor Complexes

Experimental crystal structures of HIV-1 protease variant-inhibitor complexes were obtained from the Protein Data Bank (PDB) where available. Structures unavailable experimentally were generated using structural alignment and homology modeling. All complexes were subject to standardized preparation procedures including assignment of protonation states, optimization of hydrogen-bonding networks, addition of missing atoms, and conversion to AMBER-compatible topology formats. Crystal waters observed in the corresponding experimental structures were retained throughout system preparation. A detailed description of structural modeling, alignment procedures, and protein preparation protocols is provided in Supporting Information Section S1.

### MD System Preparation

Protein and inhibitor force field parameters were generated using the ff19SB and GAFF2 force fields, respectively. The inhibitor structures were optimized using the Hartree-Fock method with the 6-31* basis set and partial charges were calculated using the Antechamber AM1-BCC method. Complexes were solvated in explicit OPC water and ionized to reproduce physiological ionic strength. Following minimization and equilibration, each protease-inhibitor complex was simulated using three independent MD replicates with randomized initial velocities. Production simulations were performed for 200 ns per replicate after equilibration, yielding a total sampling time of 600 ns per complex. Complete details of inhibitor parameterization and system construction are provided in Supporting Information Sections S2-S3.

### MD Simulations

MD simulations were performed using Amber 22 software^69^ with the PMEMD engine accelerated on a variety of NVIDIA GPUs on the Purdue University Anvil supercomputer. Each HIV-1 protease-inhibitor complex was subjected to a multistage minimization, heating, and equilibration protocol prior to production simulation. Following equilibration, three independent production trajectories were generated for each complex using randomized initial velocities to improve conformational sampling.

Production simulations were performed under periodic boundary conditions at 298K and 1 bar using the Langevin thermostat and the Berendsen barostat. Long-range electrostatic interactions were treated using particle-mesh Ewald simulation, and covalent bonds involving hydrogen atoms were constrained using SHAKE. A nonbonded cutoff of 12 Å was applied for short-range vdW and electrostatic interactions. After discarding the equilibration period, 200 ns from each replicate trajectory were retained for analysis, resulting in 600 ns of aggregate sampling for each protease-inhibitor complex. Detailed equilibration and production simulation protocols are provided in the accompanying GitHub repository (https://github.com/SchifferLab/ROBUST-2026).

### Extraction of Energetic Features from MD Trajectories

Coordinates from production trajectories were analyzed using CPPTRAJ. To characterize the energetic interactions governing inhibitor potency, pairwise residue-residue and residue-inhibitor interaction energies were calculated throughout the equilibrated simulations.

Trajectory snapshots were sampled at 20 ps intervals, and pairwise vdW and ES interaction energies were computed for all atom pairs. Atomic interaction energies were subsequently aggregated to the residue-residue and residue-inhibitor levels using custom Python scripts. These aggregated energetic descriptors were selected because resistance-associated changes in HIV-1 protease dynamics manifest through redistribution of energetic interactions throughout the structural ensemble.

For each complex, the resulting feature set consisted of 1,503 vdW interactions and 985 ES interactions. Together, these descriptors formed a physics-based representation of the protease-interaction network that served as input for statistical learning.

### Conversion of Experimental Potencies to Binding Free Energies

Experimental inhibitor constants (*K_i_*) were converted to standard free energies of binding using:

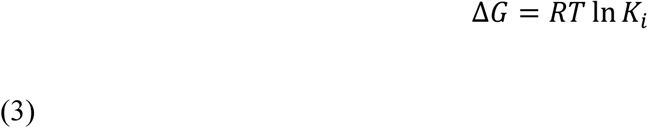

where R is the gas constant (0.001987 kcal mol^−1^ K^−1^), *T* is the temperature (298 K), and *K_i_* is expressed in molar units.

Binding free energies were used as the dependent variable for all subsequent modeling and statistical analyses.

### Identification of Recurring Energetic Descriptors

To identify energetic interactions associated with inhibitor potency, elastic net regression was applied to the MD-derived feature set using Scikit-learn. Elastic net regularization was selected because it performs feature selection while retraining correlated predictors, thereby enabling identification of recurring energetic networks rather than isolated interactions. Training datasets were repeatedly subsampled using 100 randomized train/test partitions, and elastic net models were independently fit to each partition using an L1 ratio of 0.75.

For each split, nonzero regression coefficients were identified and ranked according to their absolute magnitudes. Features that consistently reappeared across independent partitions were considered recurring descriptors of potency. This procedure reduced the original high-dimensional energetic feature space to a compact set of interpretable interactions suitable for downstream model construction and analysis.

### Generation and Selection of Predictive Models

Predictive models were constructed from the recurring energetic descriptors identified by elastic net regression. OLS regression models were generated from all possible combinations of the top-ranked descriptors containing between three and eight features. All features were standardized prior to model fitting.

Model quality was assessed using AIC, which balances goodness of fit against model complexity. Relative likelihoods were calculated from the AIC values of all candidate models, and models with relative likelihood greater than 0.05 were considered statistically supported. Feature recurrence within this ensemble of supported models was subsequently analyzed to identify robust energetic signatures associated with inhibitor potency. To determine optimal model complexity, predictive performance was evaluated as a function of feature number. The selected feature count corresponded to the point at which additional descriptors produced minimal improvement in model performance, indicating convergence of predictive accuracy.

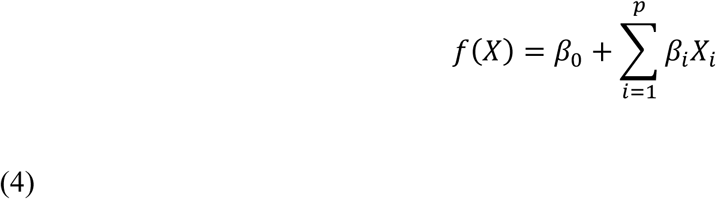

where *X_i_* represents a selected energetic descriptor, *β_i_* is its regression coefficient, and *β*_0_ is the intercept term.

### Evaluation of Model Performance and Robustness

Model performance was evaluated using both training and independent test datasets. Predictive accuracy was quantified using root-mean-square error (RMSE), Pearson correlation coefficient (*r*), Spearman rank correlation coefficient (ρ), and coefficient of determination (*R*^2^). To assess the robustness of model selection and predictive performance, the complete modeling workflow was repeated across 100 randomized train/test partitions.

Performance statistics, feature recurrence, and model-complexity convergence were summarized across the resulting ensemble of supported models. Generalization beyond the training chemical space was evaluated through independent holdout experiments in which inhibitors containing selected P1, P1′, or P2′ chemotypes were excluded from model training and reserved for external validation.

The evaluation metrics were calculated according to:

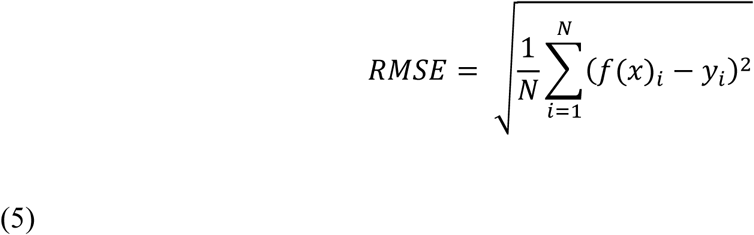

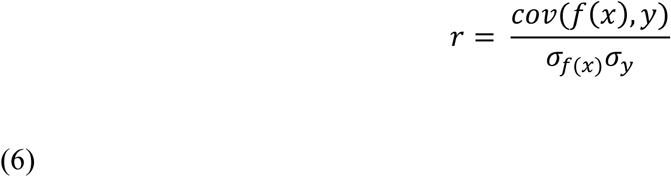

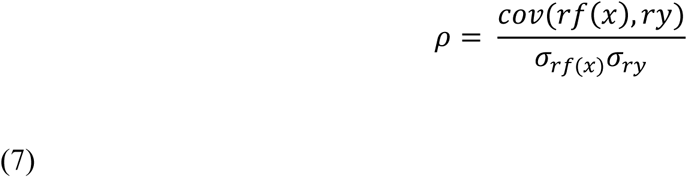

where N is the number of observations, *f*(*x*) is the model prediction, and *y* is the experimentally measured binding free energy.

### Evaluation on Unseen Chemical Modifications

To assess model generalization beyond the chemistry used for model development, independent train/test partitions were generated based on inhibitor substructure identity. Three holdout schemes were constructed corresponding to P1, P1′, or P2′ substituent classes.

For each scheme, inhibitors containing the designated chemotype were excluded from model training and reserved exclusively for external validation. All feature selection, model generation, and parameter optimization were performed using only the corresponding training set. Predictive performance of the resulting models was subsequently evaluated on the held-out compounds to determine whether the energetic signatures generalized to previously unseen inhibitor modifications.

### Ranking of Unseen Inhibitors

To evaluate the utility of ROBUST for resistance-aware lead optimization, inhibitor potencies predicted by the holdout models were averaged across the panel of HIV-1 protease variants. Inhibitors were subsequently ranked according to their predicted mean binding free energies. Predicted rankings were compared with rankings obtained from experimentally determined binding free energies to assess the ability of the energetic signatures to prioritize compounds with broad potency against resistant variants. The resulting ranking analysis provided a practical evaluation of the framework’s utility for lead optimization beyond prediction of individual binding affinities.

## 7. DATA AND CODE AVAILABILITY

All scripts used for MD trajectory analysis, energetic feature generation, statistical learning, model selection, and figure preparation are available through the accompanying GitHub repository (https://github.com/SchifferLab/ROBUST-2026). Input structures, simulation protocols, processed datasets, and analysis workflows required to reproduce the reported analyses are provided in the repository and Supporting Information.

## ASSOCIATED CONTENT

### Supporting Information

The following files are available free of charge. Supplemental Tables and Figures (PDF)

## AUTHOR INFORMATION

### Author Contributions

The manuscript was written through contributions of all authors. All authors have given approval to the final version of the manuscript.

## ACKNOWLEDGMENT

This study was supported by NIH National Institute of General Medical Sciences grants R01 GM135919 and R35 GM151996, and NSF ACCESS Allocations Projects BIO250064 and BIO260054 for GPU time.

## ABBREVIATIONS

HIV: human immunodeficiency virus
MD: molecular dynamics
ML: machine learning
vdW: van der Waals
ES: electrostatic
FEP: free energy perturbation
RMSE: root-mean-square error
LIG: ligand
AIC: Akaike Information Criterion
PDB: protein data bank.

