## Supporting Information PDF for "A Dynamics-Informed Machine Learning Framework for Inhibitor Optimization Against Quickly Evolving Targets to Avoid Resistance"

**S1. Structural Modeling and Preparation of HIV-1 Protease-Inhibitor Complexes**

X-ray crystallographic structures of the 28 inhibitors bound to HIV-1 protease Var01, as well as DRV bound to Var02, Var03, and Var04 were obtained from the PDB. To obtain the starting structures of Var02, Var03, and Var04 in complex with the remaining 27 inhibitors, PDB structures of the three variants were each aligned with the structure of the inhibitors bound to I84V variant in PyMol on residues 24, 25, 26, 85, 86, 87, and 88 from both the A and B chains. The DRV and I84V structures were removed from the alignment, and the remaining variant and inhibitor were exported as a PDB file. The remaining complexes containing variants Var05 through Var13 with inhibitors mol02 through mol28 were homology modeled starting from the crystal structure of I84V protease using Schrodinger Prime[1,2]. During homology modeling, all crystallographic waters were maintained, and the predicted structures were briefly minimized using the OPLS3 force field to resolve steric clashes.

All HIV-1 protease variant-inhibitor structures were prepared for MD simulation following the same procedure. First, PlayMolecule ProteinPrepare[3] was used to titrate protonation states, optimize the hydrogen bond network, and add missing atoms. In all complexes, the catalytic aspartate of chain A was protonated. Then, a custom script was run on the output from PlayMolecule to remove $SO_4^-$ ions from the structures and order the complex by protein, inhibitor, crystallographic waters. Finally, pdb4amber was applied to the complex to renumber the residues and rename atoms in AMBER compatible format. To ensure consistency, the chain identifications of the two monomers were exchanged if necessary: in all structures, chain B was in contact with the aniline moiety of DRV, while chain A was in contact with the bis-THF moiety.

### S2. Inhibitor Parameterization

Mol2 files were generated from the PDB structures of I84V bound to each of the 28 inhibitors. In a custom Juypter notebook, the 28 inhibitor structures from the mol2 files were aligned and renumbered according to the common minimum substructure using RDKit. Using openbabel[4], input files for Quantum Mechanical (QM) geometry optimization in Gaussian 16[5] were generated for each inhibitor. The inhibitor structures were optimized using the Hartree-Fock method with the 6-31* basis set.

Partial charges were calculated from the QM optimized structure using Antechamber AM1-BCC[6] charge calculations. The inhibitor force field parameters were obtained from GAFF2[7] using parmchk2. Finally, tleap was used to generate the library file of force field parameters for each inhibitor.

### S3. Molecular Dynamics System Construction

Tleap was used to generate the system topology and parameter files for each HIV-1 protease variant-inhibitor complex. Each system was solvated with the OPC water model[8] in a truncated octahedron, ensuring at least 10 Å between any solute atom and the periodic boundaries. Counter ions ($Na^+$ or $Cl^-$) were added by tleap to neutralize the system's charge. The volume of the resulting system was used to calculate the number of buffer ions needed to achieve a physiological concentration of 0.15 M excess salt, which resulted in a total of 35 $Cl^-$ and 35 $Na^+$ ions. AMBER force field parameters from ff19SB[9] were used from protein atoms. The resulting parameter and topology files were used as input for the MD simulations.

**Table S1.** Experimentally determined Ki values with standard errors. Newly determined Ki values are in bold, and those previously published are highlighted in yellow[10] or blue[11].

| | Var01 | Var02 | Var03 | Var04 | Var05 | Var06 | Var07 | Var08 | Var09 | Var10 | Var11 | Var12 | Var13 |
|---|---|---|---|---|---|---|---|---|---|---|---|---|---|
| **DRV** | 25.6 ± 5.6 pM | 12.1 ± 0.9 nM | 3.9 ± 0.3 nM | 12.8 ± 0.4 nM | 26 ± 1.4 nM | 77.6 ± 3.1 nM | 12.1 ± 0.5 nM | 58.3 ± 2.2 nM | 11.7 ± 0.5 nM | 57.6 ± 1.2 nM | 156.4 ± 3.67 nM | 57.0 ± 1.5 nM | 759.2 ± 57.2 nM |
| **D2** | 1.9 ± 0.7 pM | **2.47 ± 0.50 nM** | **4.79 ± 0.22 nM** | 0.84 ± 0.06 nM | **0.61 ± 0.06 nM** | **5.14 ± 0.32 nM** | **2.8 ± 0.2 nM** | **8.9 ± 0.5 nM** | **9.9 ± 0.3 nM** | **33.5 ± 2.3 nM** | 1.69 ± 0.1 nM | **41.8 ± 1.2 nM** | **42.8 ± 5.8 nM** |
| **D3** | 27.34 ± 3.85 pM | **9.46 ± 1.33 nM** | **7.79 ± 0.85 nM** | 20.79 ± 1.5 nM | **52.2 ± 0.91 nM** | **48.58 ± 6.79 nM** | **76.8 ± 4.1 nM** | **165 ± 9.1 nM** | **94.0 ± 8.1 nM** | **98.0 ± 9.3 nM** | 186.2 ± 7.5 nM | **244.0 ± 10.6 nM** | **792.5 ± 59.4 nM** |
| **D4** | 4.3 ± 0.9 pM | **3.66 ± 1.37 nM** | **4.67 ± 0.39 nM** | 1.2 ± 0.04 nM | **1.2 ± 0.12 nM** | **14.25 ± 2.43 nM** | **2.5 ± 0.2 nM** | **18.4 ± 1.0 nM** | **8.9 ± 0.5 nM** | **36.0 ± 3.0 nM** | 2.27 ± 0.3 nM | **103.4 ± 9.2 nM** | 21.5 ± 3.8 nM |
| **D5** | 2.9 ± 0.8 pM | **0.59 ± 0.29 nM** | **1.07 ± 0.10 nM** | 0.33 ± 0.07 nM | **0.17 ± 0.01 nM** | **6.89 ± 0.99 nM** | **1.2 ± 0.1 nM** | **5.0 ± 0.4 nM** | **5.8 ± 0.5 nM** | **13.0 ± 1.1 nM** | 0.44 ± 0.034 nM | **48.9 ± 2.0 nM** | **17.7 ± 2.5 nM** |
| **U1** | 26.1 ± 3.7 pM | 11.1 ± 0.5 nM | 17.8 ± 2.9 nM | 34.83 ± 2.33 nM | 29.7 ± 2.0 nM | 42.2 ± 1.8 nM | 34.6 ± 1.2 nM | 99.7 ± 5.5 nM | 23.4 ± 0.9 nM | 54.7 ± 2.8 nM | 199.1 ± 13.4 nM | 76.46 ± 4.57 nM | 1264 ± 103 nM |
| **U2** | 3.1 ± 1.0 pM | 0.93 ± 0.1 nM | 1.4 ± 0.07 nM | 4.55 ± 0.5 nM | 2.4 ± 0.22 nM | 5.1 ± 0.2 nM | 7.2 ± 0.1 nM | 12.1 ± 0.5 nM | 30.4 ± 1.2 nM | 24 ± 0.7 nM | 8.8 ± 0.7 nM | 194.3 ± 11.3 nM | 38.5 ± 2.7 nM |
| **U3** | 9.9 ± 2.7 pM | 5.3 ± 0.1 nM | 2.6 ± 0.08 nM | 35.4 ± 3.7 nM | 29.3 ± 4.1 nM | 17.9 ± 0.5 nM | 108.1 ± 5.1 nM | 103.1 ± 5.5 nM | 120.4 ± 1.2 nM | 54.9 ± 2.5 nM | 140.7 ± 4.7 nM | 175.6 ± 7.9 nM | 504 ± 29 nM |
| **U4** | 10.5 ± 1.8 pM | 0.81 ± 0.08 nM | 0.8 ± 0.03 nM | 7.3 ± 0.2 nM | 3.1 ± 0.4 nM | 5.1 ± 0.2 nM | 5.4 ± 0.3 nM | 39.7 ± 2.3 nM | 26.8 ± 1.3 nM | 30.0 ± 1.9 nM | 21.3 ± 1.7 nM | 240.6 ± 13.6 nM | 50.8 ± 2.7 nM |
| **U5** | 7 ± 1.7 pM | 1.1 ± 0.2 nM | 0.2 ± 0.08 nM | 1.89 ± 0.2 nM | 1.0 ± 0.13 nM | 1.2 ± 0.06 nM | 3.1 ± 0.1 nM | 14.0 ± 0.8 nM | 17.9 ± 0.8 nM | 17.1 ± 0.4 nM | 1.0 ± 0.17 nM | >1uM | 68.3 ± 3.0 nM |
| **U6** | 12.8 ± 3.1 pM | 3.0 ± 0.2 nM | 18.6 ± 1.7 nM | 14.92 ± 1.1 nM | 9.0 ± 0.25 nM | 7.1 ± 0.2 nM | 49.4 ± 2.4 nM | 24.2 ± 0.8 nM | 34.2 ± 1.6 nM | 62.2 ± 4.6 nM | 83.5 ± 3.8 nM | 17.6 ± 0.89 nM | 263.9 ± 8.9 nM |
| **U7** | 12.1 ± 4.5 pM | 4.2 ± 0.4 nM | 1.2 ± 0.1 nM | 6.3 ± 0.2 nM | 0.86 ± 0.08 nM | 0.07 ± 0.004 nM | 7.2 ± 0.2 nM | 7.4 ± 0.4 nM | 36.1 ± 2.1 nM | 25.8 ± 1.0 nM | 3.3 ± 0.2 nM | 13.3 ± 1.0 nM | 11.4 ± 0.5 nM |
| **U8** | 4.7 ± 1.8 pM | 0.73 ± 0.04 nM | 3.0 ± 0.15 nM | 7.58 ± 0.5 nM | 5.0 ± 0.3 nM | 1.9 ± 0.07 nM | 38.1 ± 2.5 nM | 16.9 ± 0.6 nM | 82.3 ± 5.4 nM | 45.8 ± 2.0 nM | 34.1 ± 1.1 nM | 19.8 ± 0.48 nM | 121 ± 3.8 nM |
| **U9** | 7.6 ± 1.6 pM | 0.45 ± 0.06 nM | 1.0 ± 0.06 nM | 6.34 ± 0.2 nM | 1.03 ± 0.10 nM | 1.2 ± 0.07 nM | 6.3 ± 0.5 nM | 10.1 ± 1.4 nM | 41.4 ± 4.3 nM | 40.9 ± 3.7 nM | 2.1 ± 0.5 nM | 43.4 ± 2.6 nM | 29.3 ± 0.9 nM |
| **U10** | 14.3 ± 9.3 pM | 0.062 ± 0.015 nM | 0.2 ± 0.1 nM | 0.82 ± 0.1 nM | 0.37 ± 0.061 nM | 0.14 ± 0.03 nM | 2.4 ± 0.1 nM | 3.7 ± 0.2 nM | 19.4 ± 0.7 nM | 22.7 ± 1.0 nM | 0.68 ± 0.1 nM | 20.4 ± 0.9 nM | 19.9 ± 2.0 nM |
| **PD2** | 2.7 ± 0.7 pM | **0.15 ± 0.03 nM** | **0.17 ± 0.02 nM** | 0.56 ± 0.048 nM | **0.46 ± 0.2 nM** | **0.82 ± 0.05 nM** | **0.8 ± 0.1 nM** | **3.4 ± 0.4 nM** | **6.3 ± 0.2 nM** | **5.1 ± 0.2 nM** | 2.1 ± 0.1 nM | **30.6 ± 1.6 nM** | 3.6 ± 0.4 nM |
| **PU1** | < 5 pM | **0.72 ± 0.06 nM** | **0.68 ± 0.06 nM** | 13.27 ± 3.49 nM | **6.3 ± 0.5 nM** | **8.8 ± 0.3 nM** | **4.9 ± 0.3 nM** | **16.7 ± 1.3 nM** | **5.9 ± 0.2 nM** | **21.51 ± 0.82 nM** | 145.8 ± 10.4 nM | **27.1 ± 1.1 nM** | 259.1 ± 16.9 nM |
| **PU2** | 1.8 ± 1.6 pM | **0.32 ± 0.03 nM** | **0.15 ± 0.01 nM** | 0.78 ± 0.066 nM | **0.5 ± 0.1 nM** | **0.35 ± 0.02 nM** | **1.0 ± 0.1 nM** | **3.4 ± 0.6 nM** | **8.9 ± 0.4 nM** | **5.8 ± 0.2 nM** | 3.7 ± 0.3 nM | **31.1 ± 1.9 nM** | 10.5 ± 0.9 nM |
| **PU3** | < 5 pM | **0.63 ± 0.02 nM** | **0.51 ± 0.05 nM** | 4.98 ± 0.40 nM | **5.5 ± 0.4 nM** | **3.9 ± 0.15 nM** | **7.4 ± 0.9 nM** | **11.2 ± 0.9 nM** | **50.4 ± 3.5 nM** | **13.57 ± 0.5 nM** | 52.85 ± 4.23 nM | **54.0 ± 2.8 nM** | 103.3 ± 3.4 nM |
| **PU4** | 5.6 ± 1.5 pM | **0.25 ± 0.04 nM** | **0.24 ± 0.03 nM** | 0.88 ± 0.13 nM | **0.55 ± 0.073 nM** | **0.84 ± 0.083 nM** | **0.8 ± 0.1 nM** | **4.9 ± 0.7 nM** | **6.6 ± 0.4 nM** | **8.6 ± 0.41 nM** | 4.2 ± 0.3 nM | **66.4 ± 2.8 nM** | 11.0 ± 1.0 nM |
| **PU5** | < 5 pM | **0.14 ± 0.03 nM** | **0.40 ± 0.11 nM** | 1.8 ± 0.5 nM | **0.04 ± 0.02 nM** | **2.70 ± 0.06 nM** | **0.7 ± 0.1 nM** | **0.5 ± 0.1 nM** | **5.2 ± 0.5 nM** | **3.6 ± 0.4 nM** | 1.2 ± 0.4 nM | **3.7 ± 0.2 nM** | 9.03 ± 4.21 nM |
| **PU6** | < 5 pM | **0.4 ± 0.02 nM** | **0.50 ± 0.07 nM** | 4.71 ± 0.73 nM | **2.2 ± 0.6 nM** | **1.4 ± 0.09 nM** | **3.9 ± 0.5 nM** | **3.5 ± 0.3 nM** | **13.3 ± 0.4 nM** | **17.77 ± 0.5 nM** | 51.63 ± 5.50 nM | **4.4 ± 0.2 nM** | 105.9 ± 7.4 nM |
| **PU7** | < 5 pM | **0.03 ± 0.009 nM** | **0.14 ± 0.03 nM** | 0.77 ± 0.056 nM | **0.074 ± 0.04 nM** | **0.037 ± 0.007 nM** | **1.1 ± 0.3 nM** | **1.2 ± 0.1 nM** | **12.7 ± 0.4 nM** | **4.15 ± 0.3 nM** | 1.26 ± 0.077 nM | **3.3 ± 0.1 nM** | 2.2 ± 0.2 nM |
| **PU8** | < 5 pM | **0.2 ± 0.1 nM** | **0.07 ± 0.02 nM** | 2.2 ± 0.4 nM | **2.6 ± 0.4 nM** | **0.4 ± 0.1 nM** | **5.6 ± 0.8 nM** | **2.4 ± 0.1 nM** | **28.2 ± 1.2 nM** | **8.6 ± 1.4 nM** | 32.1 ± 4.3 nM | **37.9 ± 2.8 nM** | 81.2 ± 10.8 nM |
| **PU9** | 4.4 ± 1.1 pM | **0.06 ± 0.01 nM** | **0.19 ± 0.02 nM** | 0.37 ± 0.033 nM | **0.12 ± 0.07 nM** | **0.16 ± 0.05 nM** | **0.8 ± 0.2 nM** | **1.4 ± 0.2 nM** | **7.1 ± 1.2 nM** | **5.6 ± 0.2 nM** | 1.6 ± 0.06 nM | **7.3 ± 0.4 nM** | 6.9 ± 0.5 nM |
| **PU10** | < 5 pM | **0.03 ± 0.008 nM** | **0.6 ± 0.1 nM** | 0.25 ± 0.08 nM | **0.06 ± 0.03 nM** | **0.02 ± 0.007 nM** | **0.18 ± 0.09 nM** | **0.2 ± 0.03 nM** | **5.6 ± 0.6 nM** | **4.9 ± 0.2 nM** | 0.24 ± 0.09 nM | **3.8 ± 0.2 nM** | 3.2 ± 0.5 nM |
| **PD5** | < 5 pM | **0.13 ± 0.09 nM** | **1.79 ± 0.27 nM** | 0.93 ± 0.2 nM | **1.03 ± 0.18 nM** | **2.13 ± 0.29 nM** | **0.8 ± 0.1 nM** | **6.8 ± 0.9 nM** | **6.8 ± 0.7 nM** | **7.6 ± 0.6 nM** | 1.9 ± 0.3 nM | **45.5 ± 3.5 nM** | 15.0 ± 3.5 nM |
| **PD4** | < 5 pM | **0.55 ± 0.20 nM** | **1.23 ± 0.25 nM** | 2.2 ± 0.5 nM | **2.73 ± 0.35 nM** | **3.95 ± 0.60 nM** | **1.3 ± 0.2 nM** | **6.5 ± 0.5 nM** | **6.9 ± 0.7 nM** | **4.5 ± 0.3 nM** | 4.5 ± 0.8 nM | **107.1 ± 9.7 nM** | 63.5 ± 15.2 nM |

**Table S2.** Description of the 13 HIV-1 protease variants used in this study. Homology models (HM) were generated for variants lacking an experimental structure.

| Variant ID | Mutations | Starting structure PDB ID | $K_i$ average across inhibitor set | $K_i$ standard deviation of across inhibitor set |
|---|---|---|---|---|
| Var01 | I84V | 6DH0 | 0.008 | 0.01 |
| Var02 | L10F, V32I, L33F, K45I, A71V, V82I, I84V | HM | 2.13 | 3.40 |
| Var03 | L10F, M46I, I50V, F53L, L63P, G73S | HM | 2.70 | 4.74 |
| Var04 | L10F, V32I, L33F, N37D, M46I, L63P, T74S, L76V, I84V, L89M | HM | 6.58 | 12.39 |
| Var05 | I13V, G16E, V32I, L33F, K45I, M46I, V82F, I84V | 6OPV | 6.77 | 9.54 |
| Var06 | L10I, V32I, L33F, K45I, M46I, I50V, A71V, V82I, I84V | HM | 9.19 | 17.87 |
| Var07 | L10F, L33F, M46I, I47V, I50V, F53L, L63P, A71V, L76S, V82I, I85V, L89I | HM | 13.80 | 23.20 |
| Var08 | L10F, V11I, I13V, V32I, L33F, K43T, M46L, I54L, A71V, V82I, I84V, L89M, T91S, Q92R | HM | 23.16 | 25.69 |
| Var09 | L10F, I13V, L33F, K45R, M46I, I47V, I50V, F53L, I54L, I66F, A71V, T74A, L76S, V82I | HM | 25.95 | 29.03 |
| Var10 | L10F, M46I, I47V, I50V, F53L, L63P, I72V, G73S, V82I, I85V | HM | 25.98 | 38.55 |
| Var11 | I13V, G16E, V32I, L33F, K45I, M46I, A71V, L76V, V82F, I84V | 6OPY | 40.88 | 63.33 |
| Var12 | L10F, I15V, M46I, I47V, I50V, F53L, A71V, V82I, I84V, L89T | HM | 97.10 | 189.95 |
| Var13 | I13V, G16E, V32I, L33F, K45I, M46I, I54L, A71V, L76V, V82F, I84V | 6OPZ | 167.12 | 302.55 |

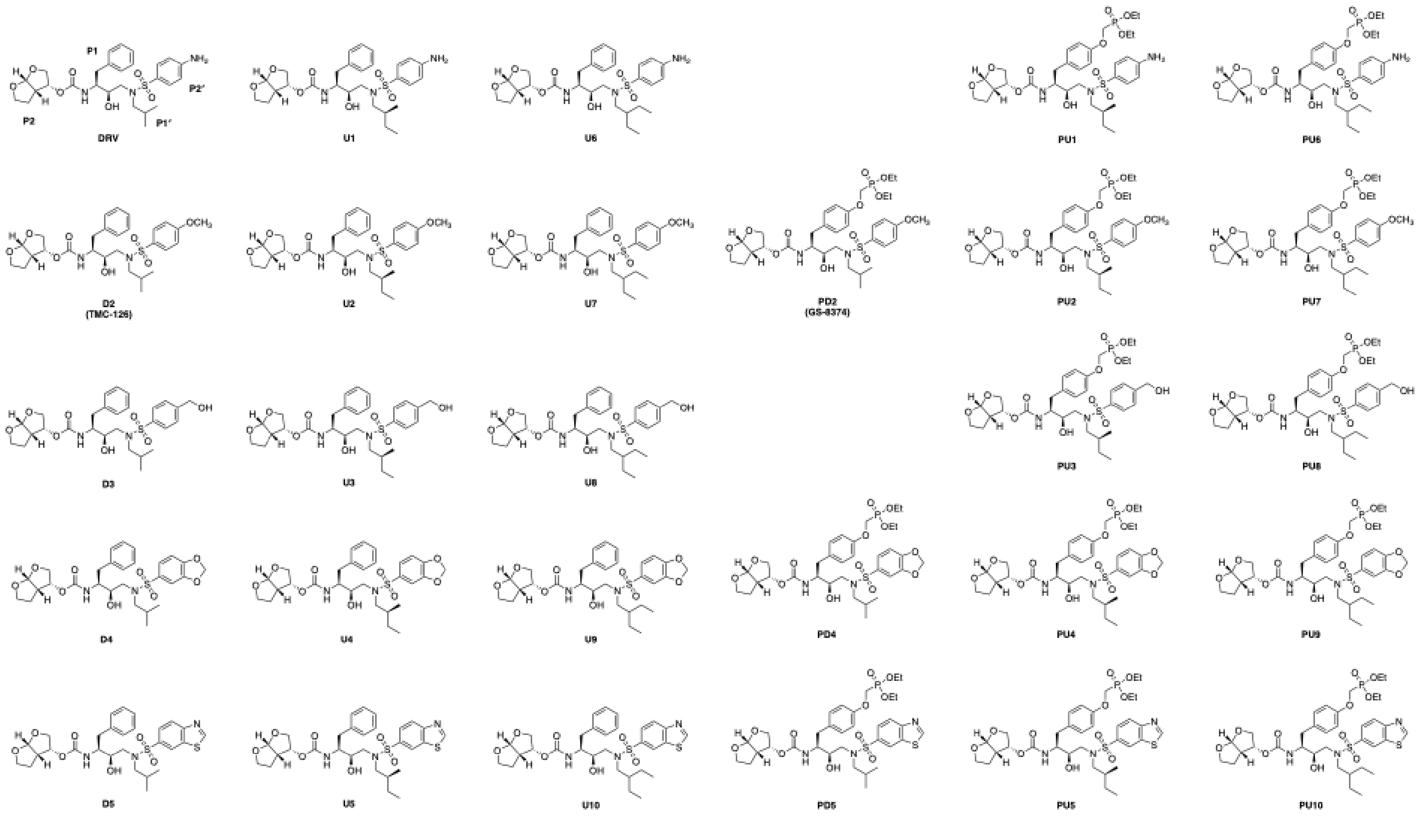


**Figure S1.** Full chemical structures of DRV and the 27 analogs used in this study.

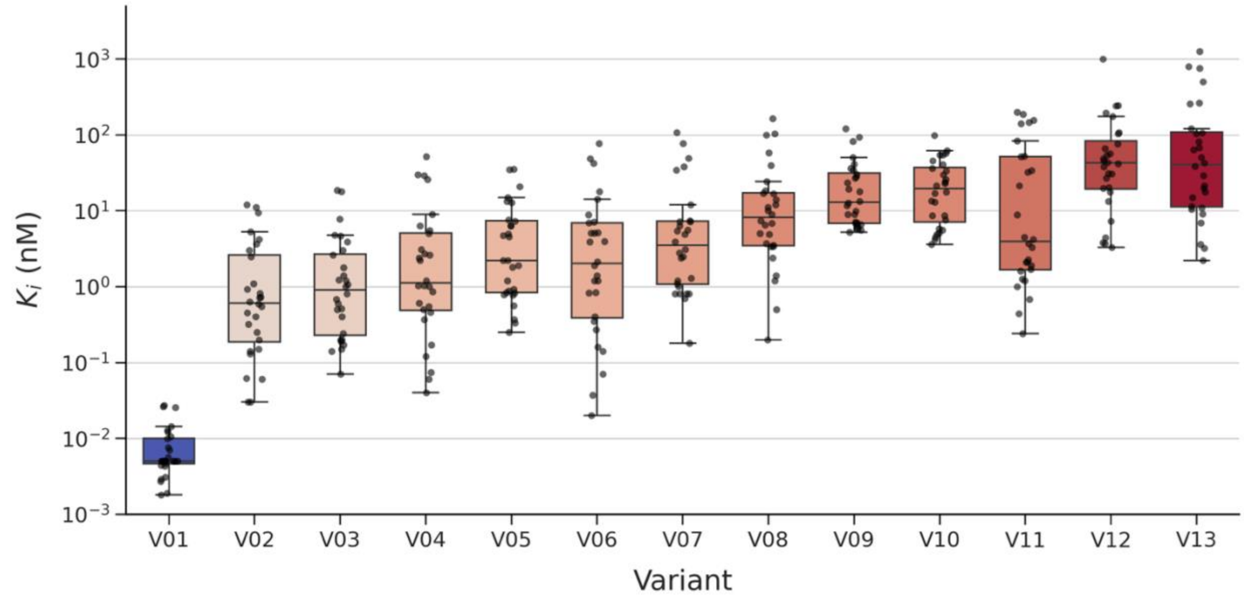


**Figure S2.** Box plot of $K_i$ values separated by variant. Variants are sorted and colored by average $K_i$ value from blue to red, where blue indicates sensitive variants and red indicates resistant variants.

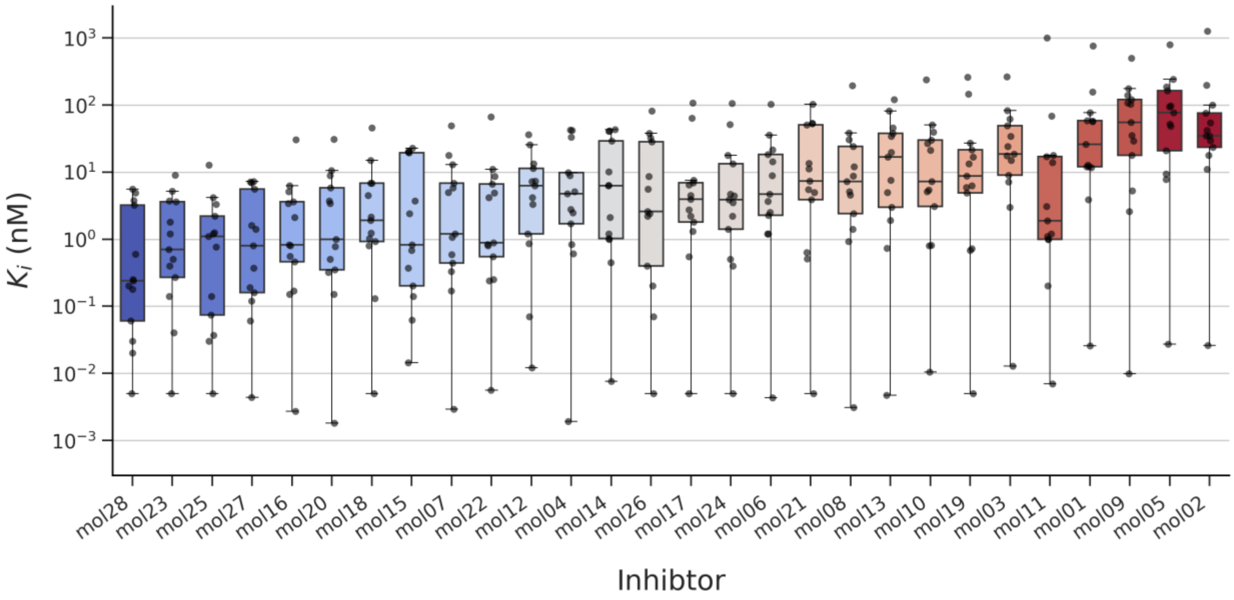


**Figure S3.** Box plot of $K_i$ values separated by inhibitor. Inhibitors are sorted and colored by average $K_i$ value from blue to red, where blue indicates higher potency and red indicates lower potency.

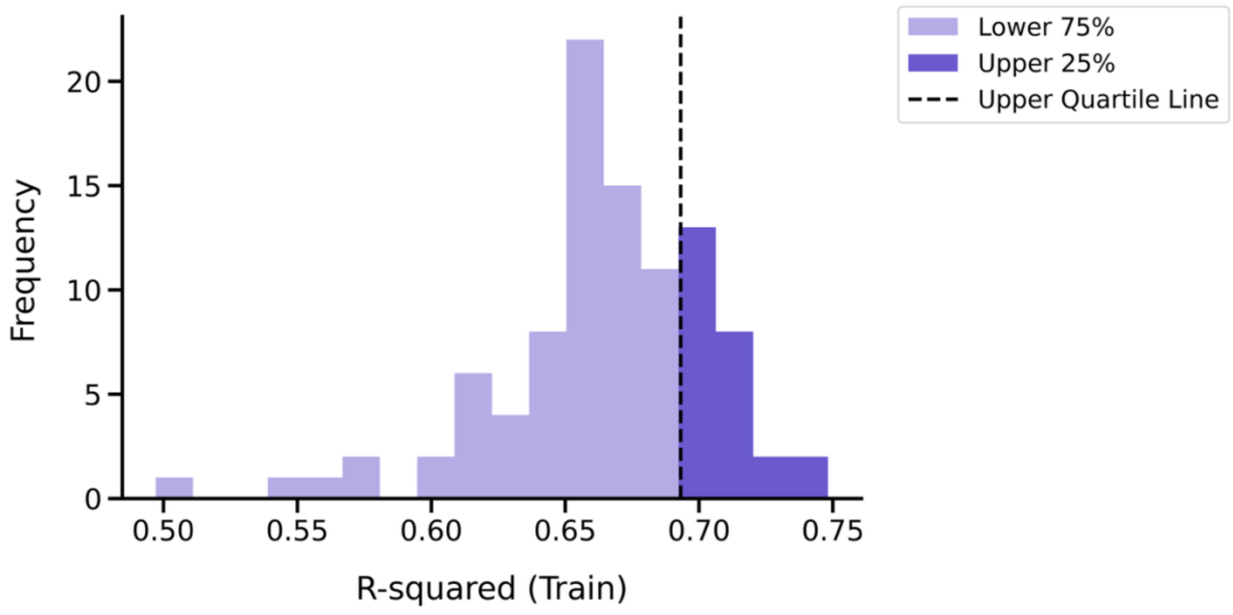


**Figure S4.** Distribution of R-squared values for the 100 models generated from random splits of the entire dataset.

A

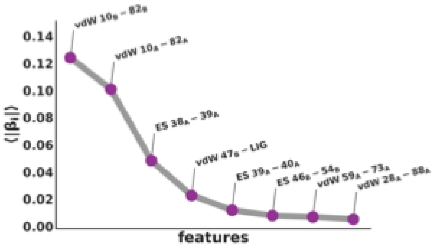


C

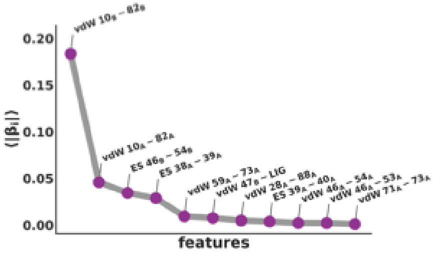


E

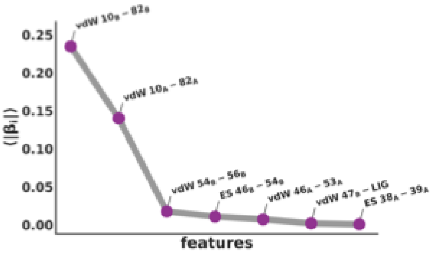


B

| Feature name | Linear Coefficient | Color (Figure 1) |
|---|---|---|
| **vdW $10_B$-$82_B$** | -0.47 | Magenta |
| **vdW $47_B$-LIG** | 0.51 | Blue |
| **ES $46_B$-$54_B$** | 0.40 | Red – Orange |
| **vdW $59_A$-$73_A$** | 0.30 | Grey |
| **vdW $28_A$-$88_A$** | -0.25 | Grey |

D

| Feature name | Linear Coefficient | Color (Figure 1) |
|---|---|---|
| **vdW $10_B$-$82_B$** | -0.47 | Magenta |
| **vdW $47_B$-LIG** | 0.47 | Blue |
| **ES $46_B$-$54_B$** | 0.47 | Red – Orange |
| **vdW $59_A$-$73_A$** | 0.28 | Grey |
| **vdW $46_A$-$54_A$** | 0.26 | Grey |

F

| Feature name | Linear Coefficient | Color (Figure 1) |
|---|---|---|
| **vdW $10_B$-$82_B$** | -0.66 | Magenta |
| **vdW $47_B$-LIG** | 0.50 | Blue |
| **ES $46_B$-$54_B$** | 0.37 | Red – Orange |
| **vdW $54_B$-$56_B$** | -0.43 | Orange – Yellow |

**Figure S5.** Ranking of physics-based features based on linear coefficients determined by elastic net regression (A, C, E) and list of features used in the final models for the inhibitor subsite-specific splits (B, D, F).

A

Random split model

| | Train | | Test | |
|---|---|---|---|---|
| # Complexes | 253 | | 109 | |
| Pearson *r* | 0.84 | | 0.80 | |
| Spearman *r* | 0.75 | | 0.73 | |
| $R^2$ | 0.71 | | 0.64 | |
| RMSE $\left(\frac{kcal}{mol}\right)$ | 0.88 | | 0.90 | |
| predictions ±1 kcal/mol | 185 | 73% | 83 | 76% |
| predictions ±2 kcal/mol | 62 | 25% | 24 | 22% |
| predictions >2 kcal/mol | 6 | 2% | 2 | 2% |

B

P1 split model

| | Train | | Test | |
|---|---|---|---|---|
| # Complexes | 195 | | 169 | |
| Pearson *r* | 0.81 | | 0.79 | |
| Spearman *r* | 0.66 | | 0.66 | |
| $R^2$ | 0.66 | | 0.48 | |
| RMSE $\left(\frac{kcal}{mol}\right)$ | 0.92 | | 1.07 | |
| predictions ±1 kcal/mol | 144 | 74% | 107 | 64% |
| predictions ± 2 kcal/mol | 48 | 25% | 51 | 31% |
| predictions > 2 kcal/mol | 3 | 1% | 9 | 5% |

C

P1' split model

| | Train | | Test | |
|---|---|---|---|---|
| # Complexes | 208 | | 154 | |
| Pearson *r* | 0.81 | | 0.84 | |
| Spearman *r* | 0.67 | | 0.76 | |
| $R^2$ | 0.65 | | 0.65 | |
| RMSE $\left(\frac{kcal}{mol}\right)$ | 0.93 | | 0.93 | |
| predictions ±1 kcal/mol | 161 | 70% | 94 | 72% |
| predictions ± 2 kcal/mol | 66 | 28% | 33 | 25% |
| predictions > 2 kcal/mol | 5 | 2% | 3 | 2% |

D

P2' split model

| | Train | | Test | |
|---|---|---|---|---|
| # Complexes | 232 | | 130 | |
| Pearson *r* | 0.84 | | 0.76 | |
| Spearman *r* | 0.72 | | 0.63 | |
| $R^2$ | 0.70 | | 0.53 | |
| RMSE $\left(\frac{kcal}{mol}\right)$ | 0.89 | | 1.01 | |
| predictions ±1 kcal/mol | 157 | 75% | 107 | 69% |
| predictions ± 2 kcal/mol | 48 | 23% | 37 | 24% |
| predictions > 2 kcal/mol | 3 | 2% | 10 | 7% |

**Figure S6.** Breakdown of accuracy in binding free energy predictions for the representative random split model (A), P1 model (B), P1' model (C), and P2' model (D).
